# Mutations in Membrin/*GOSR2* reveal stringent secretory pathway demands of dendritic growth and synaptic integrity

**DOI:** 10.1101/142679

**Authors:** Roman Praschberger, Simon A. Lowe, Nancy T. Malintan, Henry Houlden, Dimitri M. Kullmann, Maria M. Usowicz, Shyam S. Krishnakumar, James J.L. Hodge, James E. Rothman, James E.C. Jepson

**Author notes:** These authors contributed equally to this work (Simon A. Lowe, Nancy T. Malintan).

## Abstract

Mutations in the Golgi SNARE protein Membrin (encoded by the *GOSR2* gene) cause progressive myoclonus epilepsy (PME). Membrin is a ubiquitously important protein mediating ER-to-Golgi membrane fusion, and hence it is unclear how these mutations result in a disorder restricted to the nervous system. Here we use a multi-layered strategy to elucidate the consequences of Membrin mutations from protein to neuron. We show that the pathogenic mutations cause partial reductions in SNARE-mediated membrane fusion. Importantly, these alterations were sufficient to profoundly impair dendritic growth in novel *Drosophila* models of *GOSR2*-PME. We also observed axonal trafficking abnormalities in this model, as well as synaptic malformations, trans-synaptic instability and hyperactive synaptic transmission. Our study highlights how dendritic growth is vulnerable even to subtle secretory pathway deficits, uncovers a previously uncharacterized role for Membrin in synaptic function, and provides a comprehensive explanatory basis for genotype-phenotype relationships in *GOSR2*-PME.

## Introduction

Secreted, membrane, endo- and lysosomal proteins are deposited into the ER after ribosomal synthesis. Subsequently, these proteins exit the ER, transition through the Golgi apparatus and reach their ultimate target sites via the trans-Golgi network – a biosynthetic route termed the secretory pathway (Palade, 1975). The transport of proteins along this path is facilitated by membrane-enclosed vesicles, and their fusion event at the cis-Golgi is mediated by the target (t−) SNARE proteins Membrin (also known as GS27; encoded by the *GOSR2* gene), Sec22b and Syntaxin-5, in concert with the vesicle (v−) SNARE Bet1 (Parlati et al., 2000; Xu et al., 2000). Similarly to other intracellular fusion steps, these proteins are necessary for fusion of opposing lipid bilayers through the formation of a quaternary SNARE complex (Hay et al., 1997; Lowe et al., 1997; Parlati et al., 2002; Volchuk et al., 2004; Xu et al., 2000). Critical to this process is the N- to C-terminal zippering along 15 mostly hydrophobic ‘layer’ amino acids (−7 to +8) within the SNARE domain of each protein (Antonin et al., 2002; Sutton et al., 1998; Zwilling et al., 2007).

Homozygous missense (G144W: layer −3) or compound heterozygous missense and deletion mutations (G144W & K164del: between layer +2 and +3) in Membrin’s SNARE motif have recently been shown to cause the severe neurological syndrome progressive myoclonus epilepsy (PME) (Corbett et al., 2011; Praschberger et al., 2015). Patients with this form of PME, termed *GOSR2*-PME, typically present with ataxia around age three, followed by cortical myoclonus and generalized tonic-clonic seizures. Despite rapid disease progression and frequent premature death, cognitive function usually remains remarkably preserved. Correspondingly, marked neurodegeneration as an underlying primary cause has not been reported (Boissé Lomax et al., 2013; Corbett et al., 2011; Praschberger et al., 2015; van Egmond et al., 2015; 2014). Given the critical role of Membrin in ER-to-Golgi trafficking and its fundamental importance in every cell of the human body, it is unclear why Membrin mutations specifically result in nervous system dysfunction and do not cause symptoms in other organs. No paralogue is present in the human genome that could functionally replace Membrin in non-neuronal cells and therefore explain the primarily neuronal phenotype.

In the present study we set out to unravel the neuronal bottle-neck of *GOSR2*-PME. To do so, we investigated the disease mechanism of *GOSR2*-PME from molecule to neuron utilizing reconstituted liposome fusion assays, patient-derived fibroblasts and *Drosophila* models. We found that the pathogenic Membrin SNARE motif mutations result in a partial loss-of-function that is nonetheless sufficient to robustly reduce dendritic growth *in vivo*. Membrin mutations also resulted in pre-synaptic retraction and physiological abnormalities at motor neuron synapses. Together, our results suggest a mechanistic basis for the multifaceted neurological features of *GOSR2*-PME patients; highlight tight trafficking demands of growing dendrites; and illustrate a close-knit dependence of synaptic integrity and neurotransmitter release on cargo trafficking through the Golgi apparatus.

## Results

### *GOSR2*-PME mutations result in partial SNARE dysfunction

The locations of the PME-causing G144W and K164del mutations in Membrin’s SNARE domain suggests defective assembly of the quaternary cis-Golgi SNARE complex, likely resulting in reduced fusion of vesicular cargo carriers with this compartment. Given the technical difficulties associated with producing mammalian Golgi SNAREs and since mammalian and yeast Golgi SNAREs are functionally conserved (Varlamov et al., 2004), we tested for SNARE defects using a well-established yeast SNARE protein liposome fusion assay (McNew et al., 2000; Parlati et al., 2002). In this assay, v-SNARE containing liposomes are loaded with the fluorophores Nitro-2-1,3 benzoxadiazol-4yl-phosphatidylethanolamine (NBD-PE) and Rhodamine-PE. Both fluorophores are lipid-bound and exhibit robust Förster resonance energy transfer. Thus, NBD fluorescence is quenched in the presence of Rhodamine. When v-SNARE liposomes fuse with unlabeled t-SNARE-containing liposomes, NBD and rhodamine molecules become spatially separated, NBD is dequenched and NBD fluorescence increases, providing a measure of liposome fusion (Struck et al., 1981).

We introduced the corresponding PME linked G176W/D196del mutations into the Membrin yeast ortholog Bos1 (Figure 1A). Subsequently, we purified wild type (WT) and mutant Bos1 along with the SNARE partners Bet1, Sec22 and Sed5 (orthologous to mammalian Syntaxin-5) in *Escherichia coli*. Purified t-SNAREs Bos1, Sec22 and Sed5 were preassembled and reconstituted into acceptor liposomes and the v-SNARE Bet1 was incorporated into the fluorescent donor liposomes (Figure 1B). Both PME mutations resulted in a reduced rate and extent of fusion compared to WT (Figure 1C, D). However, these rates were significantly higher relative to a negative control where the Bos1-Sed5-Sec22 t-SNARE complex was omitted (Figure 1C, D). This finding implies that both GOSR2-PME mutations cause only a partial and not complete loss of function. The magnitude of these effects is consistent with the position and nature of each mutation within the SNARE motif. The D196del deletion likely results in misalignment of the subsequent hydrophobic layers in the C-terminal half of the SNARE domain, a region that provides the critical force to drive membrane fusion (Zhang, 2017). Correspondingly, this mutation reduced total fusion (after 120 min) by ~ 60% (Figure 1C, D). The G176W mutation also resulted in fusion impairment, but to a lesser degree than D196del (~ 30%; Figure 1C, D). This is consistent with an alteration in the N-terminal SNARE region that mediates the initial engagement of SNARE domains bridging two opposing lipid bilayers. Indeed, in accordance with a selective N-terminal assembly defect, the effects of G176W but not D196del were rescued by addition of a peptide comprising of the C-terminal half of the Bet1 SNARE domain, which acts to pre-structure the N-terminus (Figure 1E, F) (Melia et al., 2002). Increasing the pool of preassembled trans-SNARE complexes by overnight pre-incubation at 4°C also restored the fusion capacity of G176W-, but not D196del-Bos1 containing liposomes (Figure S1B, C), further demonstrating that the G176W and D196del mutations affect distinct regions of the Membrin SNARE domain. Taken together, these results suggest that the orthologous G144W and K164del mutations in Membrin partially impair distinct steps of the cis-Golgi SNARE complex formation, which is necessary for fusion of vesicular cargo-carriers with the cis-Golgi (Hay et al., 1997; 1998).

**Figure 1.**
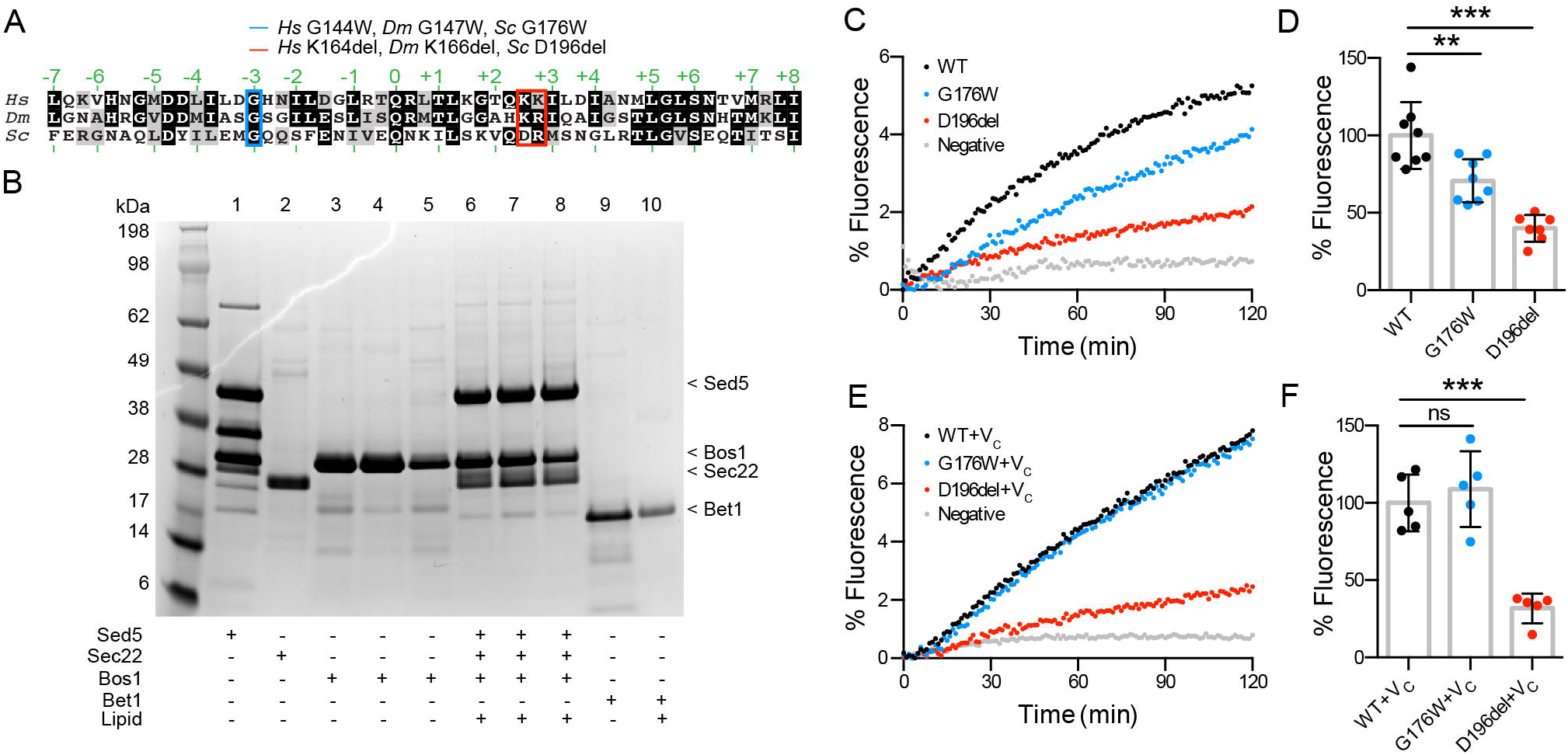
Reduced liposome fusion rates due to orthologous *GOSR2*-PME mutations. (A) SNARE domain alignment of *Homo sapiens* (*Hs*), *Drosophila melanogaster* (*Dm*) and *Saccharomyces cerevisiae* (*Sc*) Membrin (uniprot ID O14653-1), Membrin (Q9VRL2) and Bos1 (P25385) respectively. Layer amino acids critical for forming the tetrameric cis-Golgi SNARE complex are indicated in green. The disease-causing G144W and K164del (one of two consecutive lysines is deleted) and the *Drosophila* and yeast orthologous residues are highlighted in blue and red. (B) Yeast Golgi SNARE proteins Sed5 (lane 1), Sec22 (lane 2) and Bos1 WT (lane 3) and G176/D196del mutants (lane 4/5) were purified and reconstituted into acceptor liposome as t-SNARE complexes comprised of Sed5/Sec22/Bos1 (lanes 6-8 respectively). Overall stoichiometry of Sed5/Sec22/Bos1 was ~1.0x/1.0x/1.2x (see Figure S1). Yeast Golgi SNARE protein Bet1 was purified (lane 9) and reconstituted into donor liposome (lane 10) containing NBD-PE and rhodamine-PE fluorescent lipids. (C) Example traces showing increase in NBD fluorescence due to fusion between WT or G176W/D196del Bos1-containing t-SNARE complex acceptor liposomes and Bet1 donor liposomes. Negative control experiments were performed by mixing Bet1 donor liposomes with empty acceptor-liposomes lacking the t-SNARE complex. Data are expressed as a fraction of maximal NBD fluorescence after addition of detergent. (D) Endpoint (120 min) quantification of experiment as described in (C), normalized to WT. n = 8, 8, 7 for WT, G176W and D196del. (E) Example traces of experiment as in (C) with the modification that 50 μM of a peptide comprising the C-terminal half of Bet1’s SNARE domain (V_C_) was added. (F) Endpoint (120 min) quantification of experiment as described in (E), normalized to WT. n = 5. Replicate values, mean and standard deviation (SD) are shown. **, *** represent p < 0.01, 0.001, ns = not significant (p > 0.05); one-way ANOVA with Dunnett’s multiple comparison test.

### Mutant Membrin retains the capability to localize to the cis-Golgi

Only Membrin localized to the cis-Golgi will be capable of mediating deposition of ER-derived cargo into this compartment. Thus we assessed the subcellular localization of overexpressed WT and G144W/K164del mutant FLAG::Membrin in primary skin fibroblasts derived from a healthy human control. Similar to WT, both mutants successfully exited the ER and co-localized with the cis-Golgi matrix protein GPP130 (Figure S2A, B and Figure 2A, B). Previously, it was reported that G144W mutant Membrin failed to localize to the cis-Golgi in a patient derived fibroblast line (Corbett et al., 2011). We therefore re-examined these cells with an experimentally validated anti-Membrin antibody (Figure S2C, D). Membrin could clearly be detected at the cis-Golgi of G144W mutant fibroblasts and did not appear to accumulate in the ER, confirming the above overexpression results in patient cells (Figure 2C, D and Figure S2E). We note that both Golgi-localized and overall Membrin levels were reduced in the single *GOSR2*-PME patient cell line (Figure S2F-H). However, there was also substantial variability in Membrin levels between healthy control lines (Figure S2F-H). Thus, from the above data we conclude that both the G144W and K164del mutant forms of Membrin retain the intrinsic capability to localize to their cis-Golgi target compartment. This suggests that the partial SNARE domain deficiencies found in liposome fusion assays are relevant to lipid bilayer fusion rates at the cis-Golgi in living cells.

**Figure 2.**
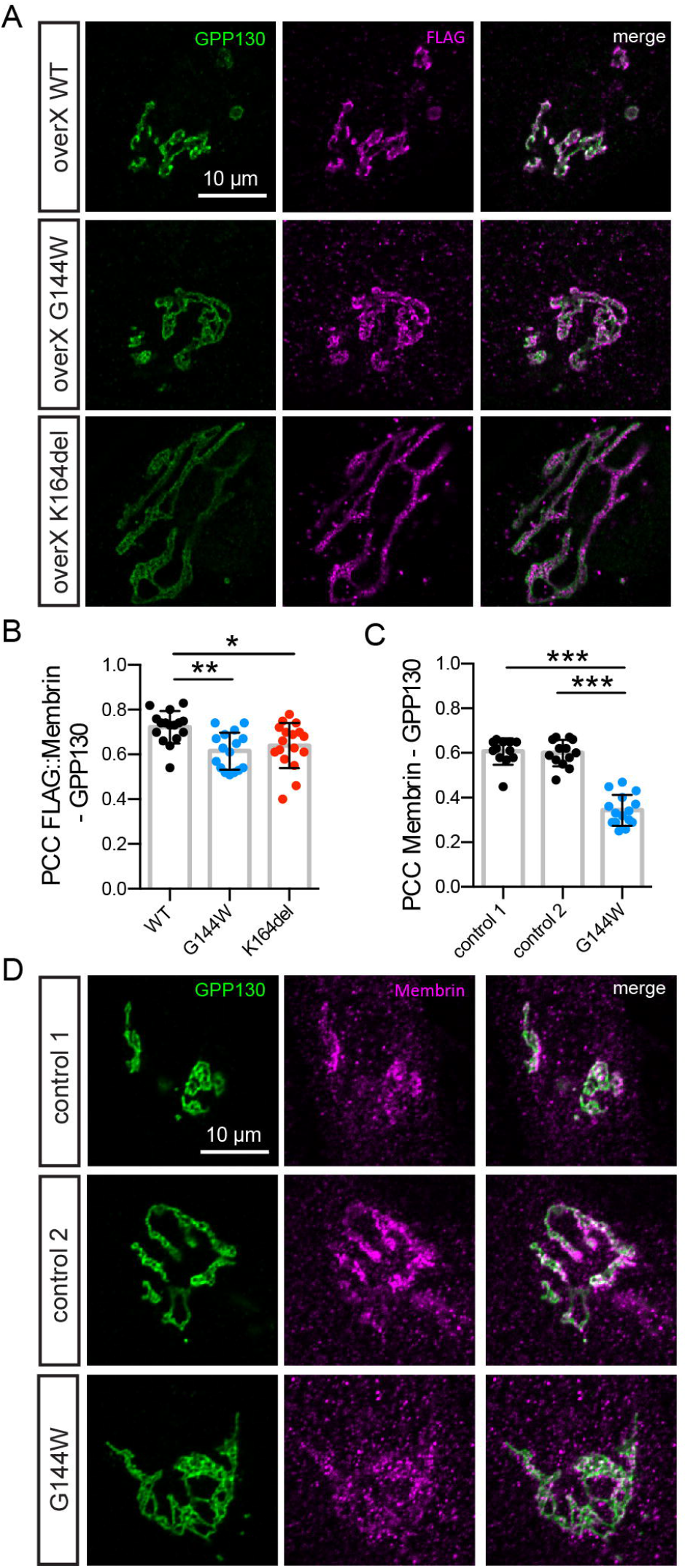
Mutant Membrin retains the capability to localize to the cis-Golgi. (A) FLAG-tagged WT and G144W/K164del mutant Membrin were overexpressed in control fibroblasts and co-stained for the FLAG tag and the cis-Golgi resident protein GPP130. Example confocal slices are shown for each overexpressed construct. (B) Pearson’s correlation coefficients between FLAG and GP130 signals of the experiment described in (A) are shown. n = 16, 16, 17 for WT, G144W and K164del. (C) Pearson’s correlation coefficients between endogenous Membrin and GP130 signals of the experiment described in (D) are shown. n = 12, 13, 15 for control 1, control 2 and G144W. (D) Example confocal slices of control and Membrin G144W mutant fibroblasts co-stained for GPP130. Replicate values, mean and SD are shown. *, **, *** represent p < 0.05, 0.01, 0.001; one-way ANOVA with Dunnett’s multiple comparison test.

### Early lethality and locomotor defects in *Drosophila* models of *GOSR2*-PME

We next sought to study the effect of Membrin mutations *in vivo* to assess their impact upon the nervous system. Golgi SNARE proteins are highly conserved throughout evolution (Kienle et al., 2009; Kloepper et al., 2007), and the *Drosophila* genome contains a single ortholog of the Membrin encoding *GOSR2* gene. In flies this gene is designated *membrin*, encoding the protein Membrin. Consistent with an essential role for Membrin orthologs in eukaryotes (Shim et al., 1991), homozygosity for the *membrin* null allele *membrin*^1524^ resulted in lethality largely prior to the L2 larval stage (Figure 3A, B) (Ghabrial et al., 2011).

**Figure 3.**
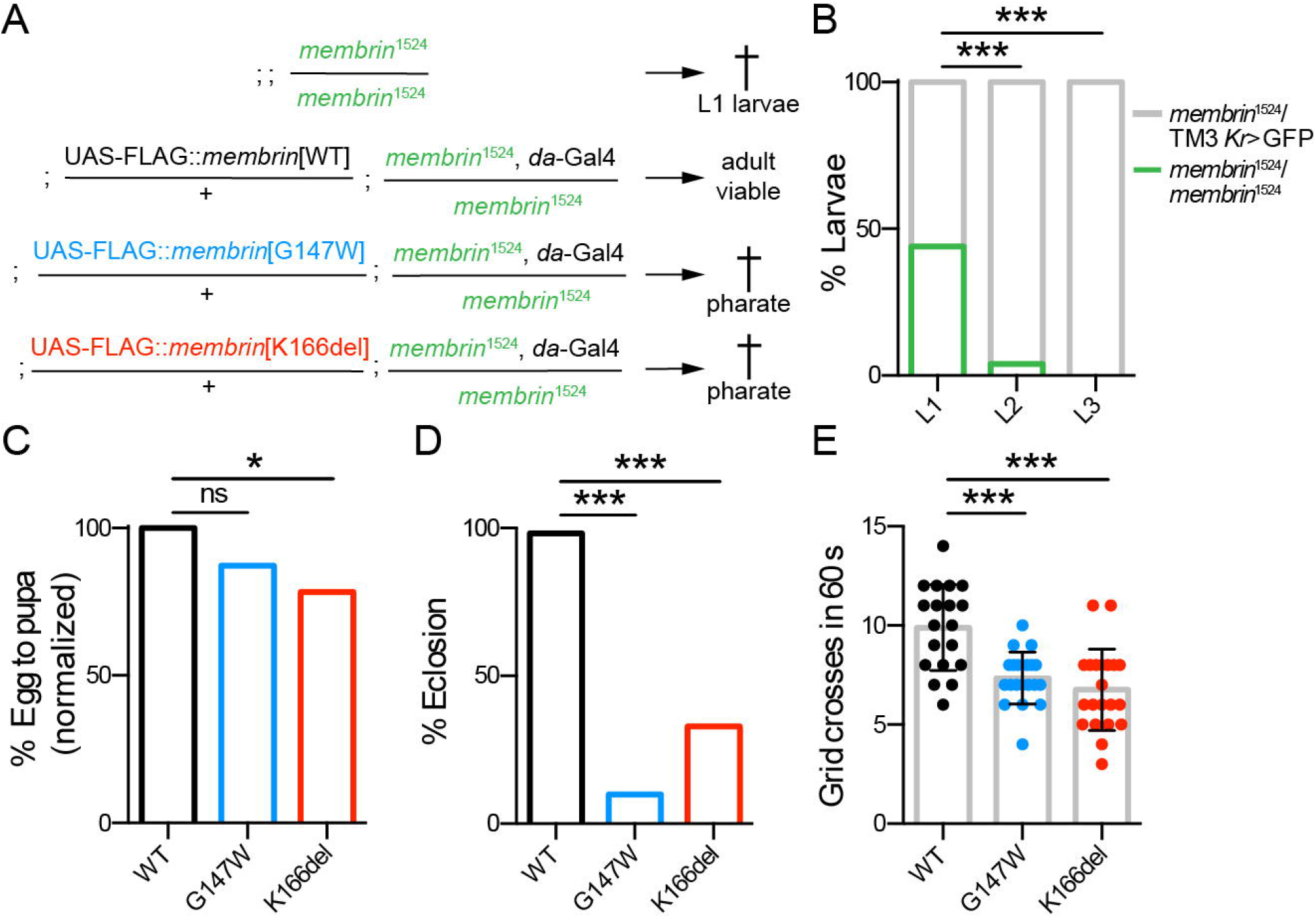
Membrin mutations cause early lethality and locomotor defects in *Drosophila*. (A) Genotypes of the *GOSR2*-PME *Drosophila* model used in this study. FLAG-tagged wild-type or mutant *membrin* (harboring the orthologous G147W/K166del mutations) is globally expressed via the *daughterless*-Gal4 driver in a *membrin* null (*membrin*^1524^) background. The shorthand Mem-WT, Mem-G147W and Mem-K166del is used throughout the paper. (B) The *membrin*^1524^ allele was balanced over the fluorescently labeled TM3 Kr>GFP chromosome to discern heterozygote animals. Homozygosity for *membrin*^1524^ caused largely L1 lethality, as at the L2 stage hardly any non GFP-positive larvae were detected. n = 50, 53, 56 for L1, L2 and L3 larvae. (C) Global expression of WT, G147W and K166del mutant Membrin rescued *membrin* null *Drosophila* to the pupal stage. Data are expressed relative to Mem-WT. n = 1222, 1308, 1260 eggs/embryos for Mem-WT/-G147W/-K166del. (D) Mem-G147W and Mem-K166del *Drosophila* frequently died as fully developed pharate adults, as shown by a drastic decrease in eclosion rates compared to Mem-WT. n = 120, 112, 97 for Mem-WT/-G147W/-K166del non-*tubby* pupae. (E) Freely moving Mem-G147W and Mem-K166del L3 larvae crossed fewer 4 mm grids in 60 s than Mem-WT. n = 19, 20, 21 for Mem-WT/-G147W/-K166del. Replicate values, mean and SD are shown. *, **, *** represent p < 0.05, 0.01, 0.001, ns = not significant (p > 0.05); Fisher’s exact test with Bonferroni correction (B-D) or one-way ANOVA with Dunnett’s multiple comparison test (E).

To assess the pathogenic *GOSR2*-PME mutations in *Drosophila*, we initially generated transgenic fly lines harboring FLAG-tagged wild-type or mutant (G144W or K164del) UAS-*GOSR2* transgenes. Each transgene sequence was integrated at the same genomic locus using site-specific ΦC31-mediated recombination in order to control for position effects on expression levels (Figure S3A) (Bischof et al., 2007). These transgenes, downstream of the yeast *upstream activating sequence* (UAS), are inert and only transcribed in the presence of the Gal4 transcriptional activator expressed via a promoter of choice (Brand and Perrimon, 1993). To enable global expression of wild-type or mutant *GOSR2* in a *membrin* null background we used the global *daughterless-Gal4* driver. Each genetic component of these models was outcrossed into an isogenic iso31 background for five generations to limit genetic variability between strains. Indeed, expression of wild-type human *GOSR2* fully rescued the lethality of *membrin* null larvae and yielded adults that appeared morphologically grossly normal (Figure S3B, C). While these animals exhibited severe motor impairments and usually died after three days, this result nonetheless demonstrates functional conservation between human and *Drosophila* Membrin, supporting the use of *Drosophila* to model *GOSR2*-PME. Because neither mutant *GOSR2* transgene rescued *membrin*^1524^ animals to the L3 larval stage (data not shown), we next generated *GOSR2*-PME models that were closer to the normal physiology of *Drosophila*. According to the identical strategy as for the human UAS-GOSR2 transgenes, we created mutant (G147W and K166del) and wild-type *Drosophila* UAS-*membrin* transgenes in order to be able to express wild-type or mutant *membrin* in a *membrin* null genetic background. For simplicity, we term these mutant fly lines and their associated control Mem-G147W, Mem-K166del and Mem-WT (Figure 3A).

In contrast to *membrin* null flies, Mem-WT flies were viable to the adult stage, indicating that transgenic FLAG-tagged *Drosophila* Membrin is functional (Figure 3C, D). Mem-G147W and Mem-K166del flies were viable to the pupal stage, surviving significantly longer than *membrin* null animals (Figure 3C). However, Mem-G147W and Mem-K166del flies frequently died within the pupal cases as fully developed pharate adults (Figure 3D). When manually released from the pupal case, Mem-G147W and Mem-K166del adults appeared weak and uncoordinated. Mutant animals that successfully freed themselves from their pupal cases usually became stuck in the fly food soon after eclosion and died within a few days. Mem-G147W and Mem-K166del L3 larvae also displayed significantly reduced rates of locomotion compared to Mem-WT (Figure 3E). Similar locomotor deficits and lack of coordination may explain the frequently observed inability of Mem-G147W and Mem-K166del flies to escape from the pupal case, leading to early lethality. Thus, although both mutants survived longer than *membrin* null flies, the incomplete nature of this rescue provides in vivo evidence that these mutations cause partial – not complete – loss of Membrin function, consistent with the above liposome fusion assays (Figure 1).

### Profound dendritic growth deficits in *GOSR2*-PME model neurons

The early-onset of neurological symptoms in *GOSR2*-PME suggests that Membrin mutations might alter aspects of neuronal development. Important clues towards the underlying mechanism stem from studies investigating neuronal consequences of mutations in other ER-to-Golgi trafficking proteins. Overexpression of GTP locked Q71L-Arf1 in developing cultured hippocampal neurons led to severely impaired dendritic growth (Horton et al., 2005). Similarly, Ye et al. found in a *Drosophila* forward genetic screen that mutations in Sec23, Sar1 and Rab1 cause dendritic growth deficiencies (Ye et al., 2007). Such early secretory pathway blocks likely prevent ER-derived lipids and proteins from reaching the plasmalemma of growing neurons, which place immense demands on such supplies due to their very large surface area (Hanus and Ehlers, 2008).

However, whereas Q71L-Arf1 and truncated Sar1 result in a complete block of anterograde trafficking (Dascher and Balch, 1994; Horton et al., 2005; Ye et al., 2007), the above liposome fusion assays and *Drosophila* GOSR2-PME models indicate that PME-causing mutations in Membrin are partial loss-of-function (Figure 1 and 3). Thus, we sought to test whether even a partial decrease in ER-to-Golgi membrane fusion was sufficient to impair dendritic growth. To do so, we genetically labeled ddaC sensory neurons within the larval body-wall with a membrane-tagged – and thus secretory pathway dependent – fluorophore (CD4::tdGFP) under control of the *ppk* promoter (Grueber et al., 2003; Han et al., 2011). These neurons have highly sophisticated, tiled dendritic arbors that branch in 2D and they are unambiguously polarized into a single axon and multiple dendrites (Figure 4A, arrow-head indicates the axon). Using this approach, we imaged the same identifiable ddaC neuron in abdominal segment 5 of Mem-WT, Mem-G147W and Mem-K166del L3 larvae (Figure 4A). Strikingly, both Mem-G147W and Mem-K166del larvae exhibited profoundly decreased dendritic length and a reduction in the number of terminal dendritic branches relative to Mem-WT (Figures 4B, C). Sholl analysis further revealed reduced elaboration of dendritic arbors and a significant reduction in total dendritic intersections in Mem-G147W and Mem-K166del (Figure 4D, E). Using fluorescence recovery after photo-bleaching (FRAP), we tested whether trafficking of the CD4::tdGFP cargo was altered in Mem-G147W and Mem-K166del dendrites. Mem-K166del larvae exhibited a clear reduction in CD4::tdGFP FRAP, and a similar albeit non-significant trend was seen for Mem-G147W (Figure 4F). Thus, dendritic growth and cargo trafficking within dendrites are impacted to a greater degree in Mem-K166del than Mem-G147W, again consistent with a more severe SNARE defect of the orthologous D196del Bos1 mutation in liposome fusion assays (Figure 1).

**Figure 4.**
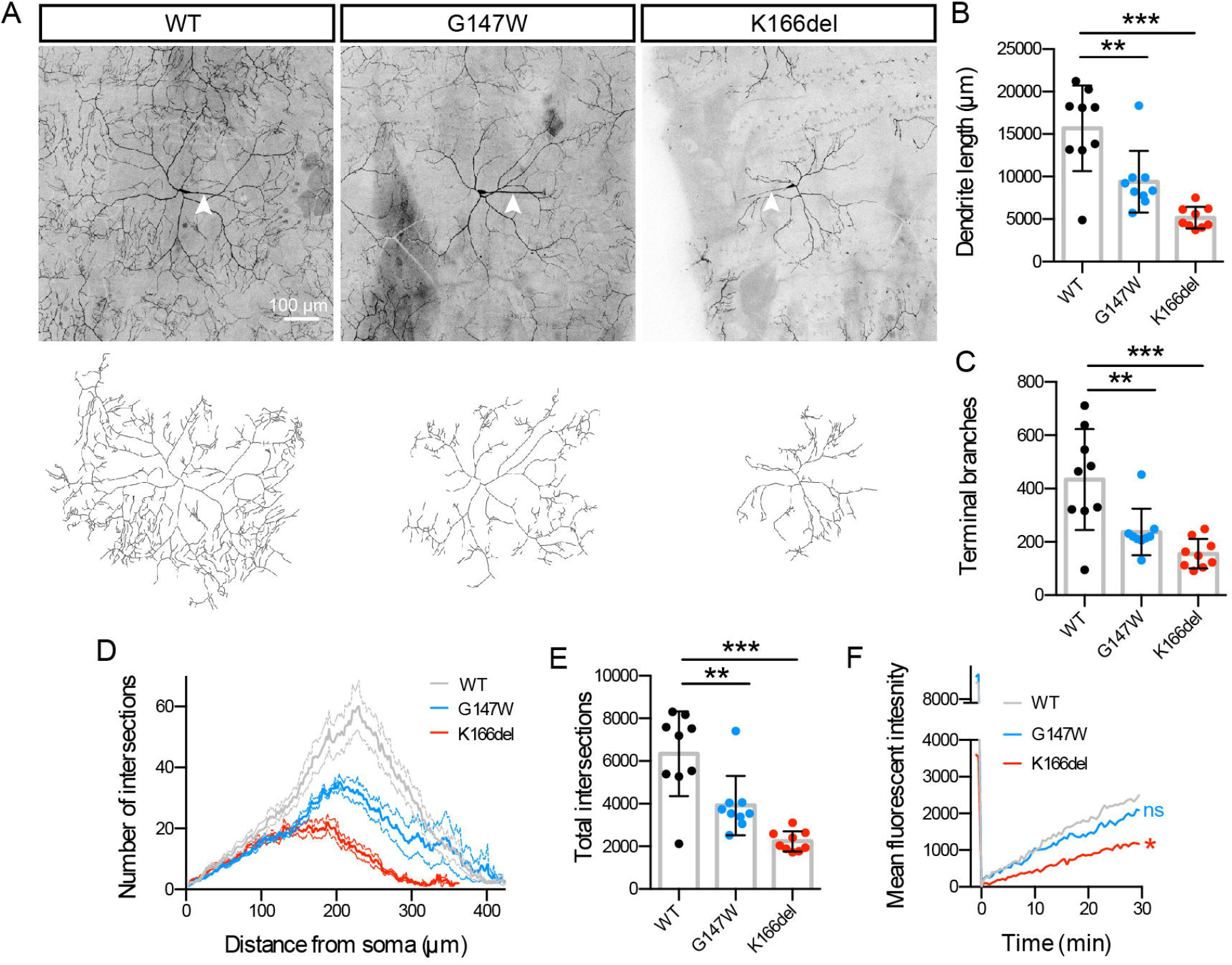
Membrin mutations cause dendritic growth deficits. (A) Maximum intensity projections of ddaC abdominal segment 5 neurons genetically labeled with *ppk* > CD4::tdGFP in Mem-WT/-G147W/-K166del. Respective tracings of the dendritic arbors are shown below. Arrowheads indicate axons. (B) Total dendritic length extracted from tracings as shown in (A). 9 A5 ddaC neurons per genotype were traced and analyzed in (B-E). (C) Number of terminal branches of ddaC A5 neurons. (D) Number of intersections of dendritic tracings with concentric circles with 2 pixel/circle increasing radii. Mean ± standard error of the mean (SEM) are shown. (E) Total intersection of Sholl analysis as shown in (D). (F) CD4::tdGFP in large segements of primary ddaC A5 dendrites adjacent to the soma were photobleached with a 50 μm^2^ region of interest and fluorescence recovery quantified 25 μm from the soma proximal bleach margin. Means of n = 9/8/9 ddaC neurons for Mem-WT/-G147W/-K166del are shown. */ns indicate endpoint comparison after 30 min recovery. Replicate values, mean and SD are shown unless otherwise stated. *, **, *** represent p < 0.05, 0.01, 0.001, ns = not significant (p > 0.05); one-way ANOVA with Dunnett’s multiple comparison test.

### Reduced cargo trafficking in Membrin mutant axons

While both growing dendrites and axons require abundant membrane addition and thus rely heavily upon secretory trafficking (Aridor and Fish, 2009), Ye et al. suggested that axonal growth can be privileged in the face of ER-to-Golgi trafficking deficits (Ye et al., 2007). We examined this notion in our *GOSR2*-PME *Drosophila* model and were able to detect axonal CD4::tdGFP signal in each ventral nerve cord (VNC) segmental nerve (Figure 5A; for saturated images see Figure S4A). This finding suggests that at least one axon derived from the three *ppk*-positive neurons per hemisegment (designated ddaC, v’ada and vdaB) reached its comparably distant target (Grueber et al., 2003; 2002; 2007). Compared with the reduced elaboration of ddaC dendrites (Figure 4), this is consistent with dendritic growth being more severely impaired than axonal growth by *GOSR2*-PME mutations. Nevertheless, axonal and pre-synaptic CD4::tdGFP levels were significantly reduced both in the VNC and in individual segmental nerves of both Mem-G147W and Mem-K166del at steady state (Figure 5A-C). Thus, a secretory pathway deficit is clearly present in distal axons of *membrin* mutant *Drosophila*. We used the large and experimentally accessible neuromuscular junction (NMJ) of L3 larvae as a model synapse to test whether the deficit in trafficking seen with CD4::tdGFP in *ppk* neurons was also reflected by changes of endogenous synaptic cargos (Harris and Littleton, 2015). Amongst several tested pre-, post- and trans-synaptic proteins, we found robust reductions in the steady state levels of the synaptic vesicle protein Cysteine String Protein (CSP), while the other cargos exhibited no consistent change compared to Mem-WT (Figure S4B-H). These results provide proof-of-principle that Membrin mutations can affect the abundance of endogenous synaptic proteins. The different effects upon individual synaptic components at steady state may reflect varying trafficking demands of synaptic proteins due to differences in synaptic turnover.

**Figure 5.**
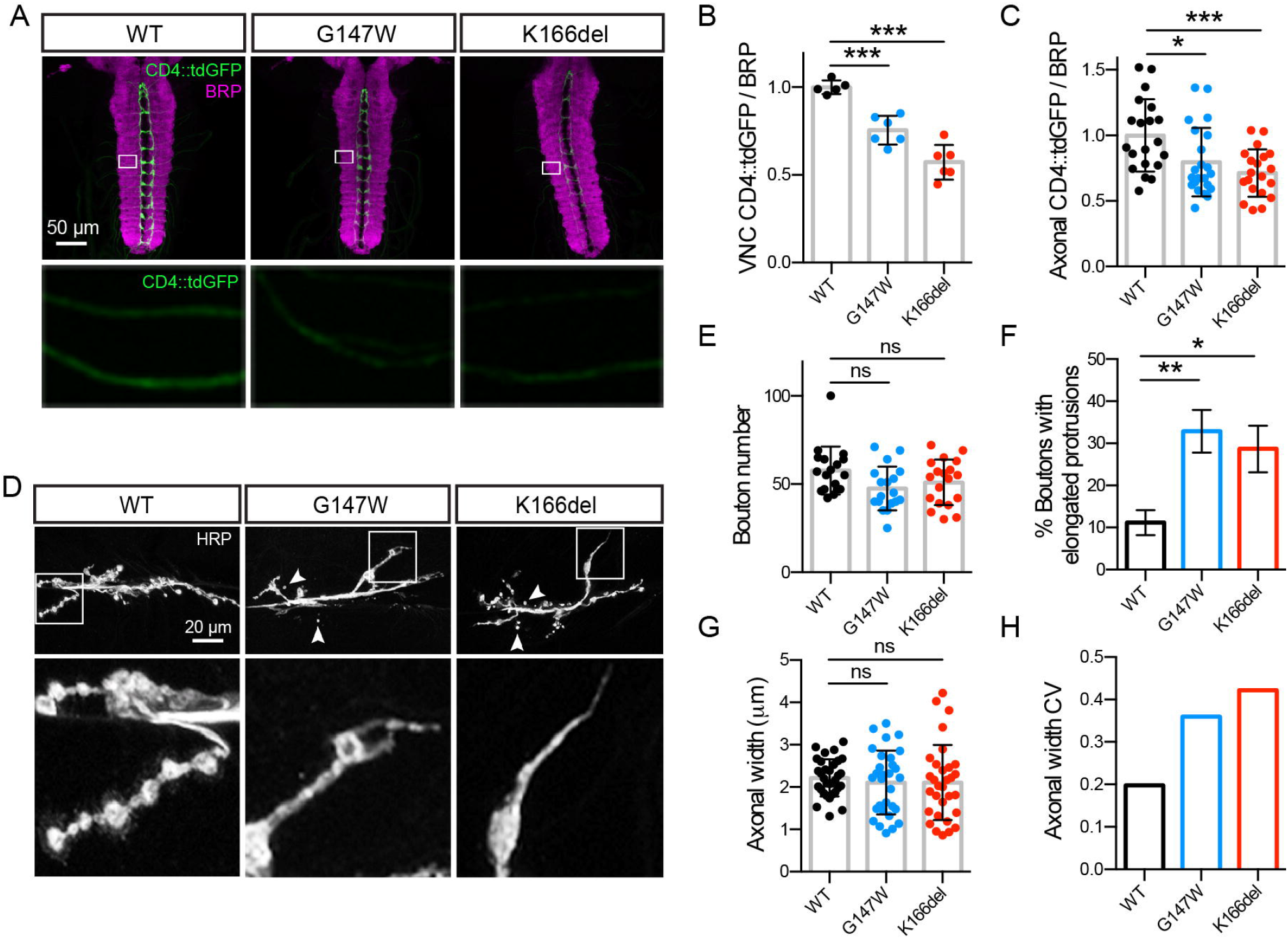
Membrin mutations alter presynaptic morphology and axonal stereotypy. (A) Top: confocal z-stacks showing projections from *ppk*-positive sensory neurons labeled with membrane-tagged CD4::tdGFP, innervating the ventral nerve cord (VNC) of L3 larvae. Synaptic neuropil of the VNC is labeled with anti-BRP. Below: magnified regions of segmental nerves. (B-C) Quantification of CD4::tdGFP fluorescence in the VNC neuropil (B) or in segmental nerves (C) normalized to BRP and expressed relative to WT. n = 5, 5, 6 (B) and 20, 22, 20 (C) for Mem-WT/-G147W/-K166del. (D) Top: example confocal z-stacks of HRP-labeled motor neurons innervating muscle 6/7, segment 3 of L3 larvae. Arrowheads point to small, isolated boutons that appear unattached to the axon. Below: magnified terminal boutons. In contrast to the rounded morphology in Mem-WT larvae, terminal boutons in Mem-G147W and Mem-K166del often exhibit elongated protrusions. Further examples are depicted in Figure S6C. (E) Average number of boutons (type 1b and 1s) at muscle 6/7, segment 3. n = 18, 18, 19 for Mem-WT/-G147W/-K166del. (F) Percentage of terminal boutons with elongated protrusions. Mean and SEM are shown. n = 32, 31, 31 for Mem-WT/-G147W/-K166del. (G) Maximal axonal width of motor neurons innervating muscle 6/7, segment 3. n = 32, 30, 31 for Mem-WT/-G147W/-K166del. (H) Coefficient of variation (calculated as the SD/mean) in axonal width for each genotype. Replicate values, mean and SD are shown unless otherwise stated. *, **, *** represent p < 0.05, 0.01, 0.001, ns = not significant (p > 0.05); one-way ANOVA with Dunnett’s multiple comparison test (B-C) or Kruskal-Wallis test with Dunn’s post-hoc test (E-G).

### Unimpaired secretory trafficking in G144W Membrin mutant fibroblasts

*GOSR2*-PME patients exhibit a restrictive neurological phenotype despite the ubiquitous expression of Membrin. We wondered whether the neuronal secretory pathway deficit observed in *GOSR2*-PME model axons might be the consequence of the unique geometry of neurons, and thus not be applicable to a non-neuronal cell. To address the possibility that the partial loss of function Membrin mutations might not be functionally relevant in a nonneuronal cell type with lower trafficking demands, we performed Golgi trafficking assays in primary skin fibroblasts from a G144W *GOSR2*-PME patient and healthy controls by overexpressing human growth hormone (hGH) fused to four FM domains and a Halo-tag. The FM domains self-aggregate in the absence of a solubilizing drug and thus do not allow ER exit of this model cargo (Figure S5A 1^st^ column). Conversely, addition of D/D solubilizer disaggregates the FM domains and allows the cargo to enter the secretory pathway. By 10 minutes after solubilization we detected significant colocalization with the cis-Golgi marker GM130, which decreased by 20 and 30 min as the cargo exited this compartment (Figure S5A, B). Remarkably, trafficking kinetics in G144W mutant Membrin fibroblasts were almost indistinguishable from controls (Figure S5A, B). This finding supports the concept that cellular effects of the partial loss of function G144W Membrin mutation are unmasked only under large secretory pathway requirements, suggesting why only the nervous system with its high trafficking demands is symptomatically affected in *GOSR2*-PME.

### Presynaptic morphological defects in *GOSR2*-PME model neurons

Because we could detect secretory pathway defects in distal axons and synapses of *GOSR2*-PME *Drosophila* models (Figure 5A-C), we next investigated whether synaptic integrity might be altered due to the *GOSR2*-PME mutations. To this end we examined presynaptic morphology at the L3 larval NMJ by labeling neuronal membranes with an anti-horseradish peroxidase (HRP) antibody, which stains neurons due to cross-reactivity with certain glycans on their surfaces (Fabini et al., 2001). These motor neuron termini are composed of an array of rounded synaptic boutons housing active zone domains responsible for coupling calcium influx to neurotransmitter release (Harris and Littleton, 2015). Motor neurons of Mem-WT, Mem-G147W and Mem-K166del all successfully formed synapses, and Mem-G147W and Mem-K166del synapses did not exhibit significant reductions in bouton number nor size relative to Mem-WT (Figure 5D, E and Figure S6A-C). However, we observed two clear morphological abnormalities in Mem-G147W and Mem-K166del synapses. Firstly, Mem-G147W and Mem-K166del terminal synaptic boutons often exhibited elongated axonal protrusions lacking rounded boutons, that were less common and shorter in Mem-WT (Figure 5D, F and Figure S6C, D). Secondly, we observed small boutons in Mem-G147W and Mem-K166del synapses that were disconnected from the main axonal branch (Figure 5D arrowheads). In addition, analysis of axonal diameter revealed a significant increase in the variability of the maximal axonal diameter (Figure 5G), as measured by the coefficient of variation (Figure 5H) and F-test (Mem-G147W vs. Mem-WT, p = 0.0035; Mem-K166del vs. Mem-WT, p = 0.0002). Thus, partial reductions in secretory pathway trafficking result in multifaceted abnormalities in motor neuron synapse development and impact the stereotypy of terminal axon morphology.

### Membrin mutations result in synaptic retraction and cytoskeletal fragmentation

To examine potential effects of Membrin mutations on synaptic structure in more detail, we co-stained Mem-WT, Mem-G147W and Mem-K166del synapses with antibodies against the presynaptic active zone marker Bruchpilot (BRP) and postsynaptic GLURIII glutamate receptors (Figure 6A). BRP localized to Mem-G147W and Mem-K166del NMJs in amounts comparable to Mem-WT (Figure 6B). However, dual pre- and postsynaptic labeling revealed pronounced strings of small presynaptic boutons in Mem-G147W and Mem-K166del synapses in which presynaptic BRP was no longer opposed to postsynaptic GLURIII (Figure 6A, C). This disruption of trans-synaptic organization is indicative of pre-synaptic retraction, where synaptic connections initially form but fail to be maintained throughout development (Eaton et al., 2002; Pielage et al., 2008).

**Figure 6.**
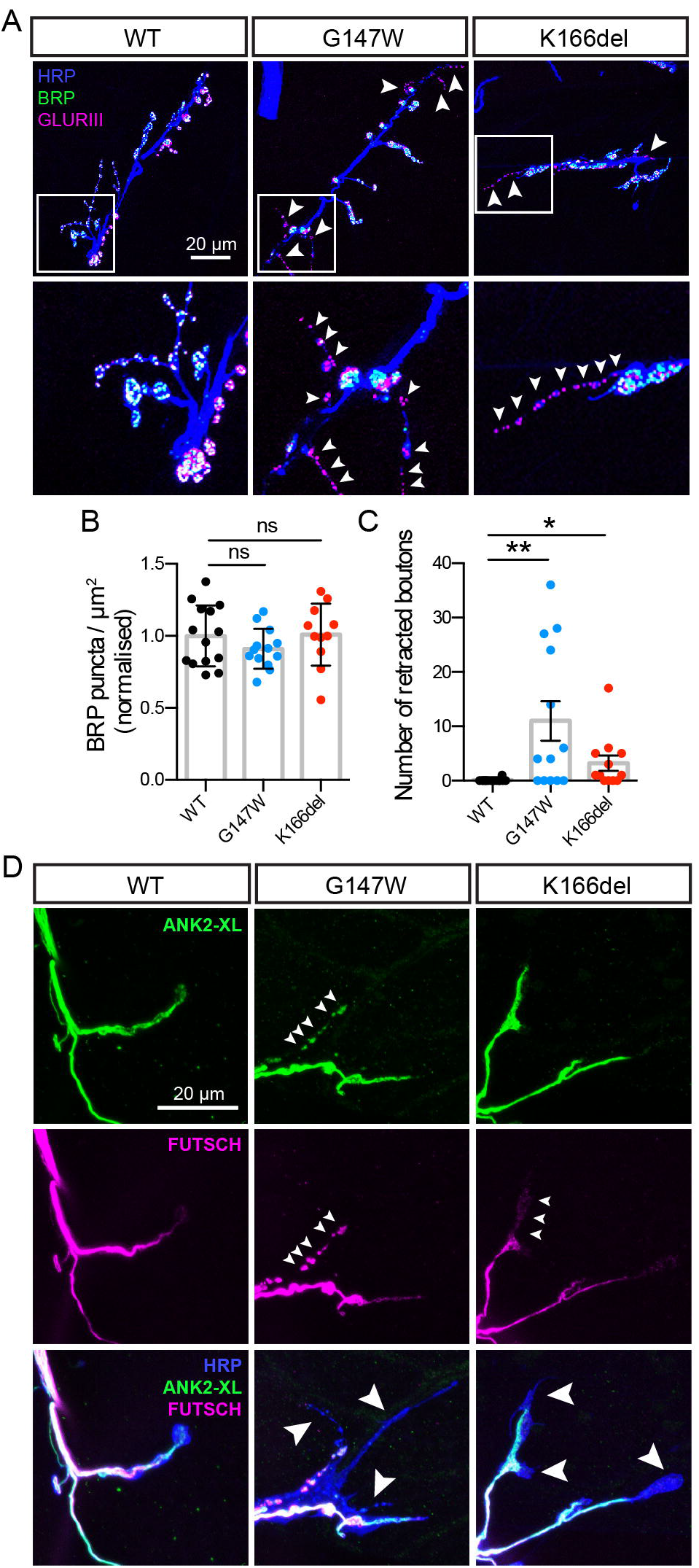
Synaptic retraction and presynaptic cytoskeletal fragmentation in *membrin* mutants. (A) Top: Maximum intensity z-projection of confocal stacks showing pre- and post-synaptic apposition between BRP-labeled active zones and post-synaptic GLURIII glutamate receptors. Arrowheads denote regions where glutamate receptors lack their presynaptic active zone counterparts. Below: magnified view of regions exhibiting loss of BRP-labeled active zones in Mem-G147W and Mem-K166del synapses. (B) Normalized density of BRP puncta per NMJ area. n = 14, 13, 11 for Mem-WT/-G147W/-K166del. (C) Average number of synaptic boutons where BRP fails to oppose GLURIII. n = 14, 13, 12 for Mem-WT/-G147W/-K166del. (D) Confocal z-stack maximum intensity projections illustrating localization of the cytoskeletal associated components Ankyrin-2-XL (ANK2-XL) and Futsch, a microtubule-binding protein. Small arrowheads point to synaptic domains containing either fragmented Futsch and ANK2-XL or reduced amounts of Futsch. Large arrowheads point to synaptic boutons and elongated protrusion apparently lacking both Futsch and ANK2-XL. Replicate values, mean and SD are shown. *,** represent p < 0.05, 0.01, ns = not significant (p > 0.05); Kruskal-Wallis test with Dunn’s post-hoc test.

Synaptic retraction can be induced by mutations in several cytoskeletal proteins (Koch et al., 2008; Pielage et al., 2008; 2011; 2005). Long protrusions lacking specialized boutons and alterations in axonal diameter in Mem-G147W and Mem-K166del synapses further suggested potential cytoskeletal defects in these backgrounds (Pielage et al., 2011; Stephan et al., 2015). To uncover molecular correlates of synaptic retraction in Mem-G147W and Mem-K166del, we therefore examined the localization of two presynaptic cytoskeleton associated proteins: Futsch (a microtubule-binding protein) and Ankyrin-2XL (ANK2-XL) (Koch et al., 2008; Roos et al., 2000). In Mem-WT synapses, both Futsch and ANK2-XL were strongly colocalized in central and distal axons and invade terminal boutons (Figure 6D) (Stephan et al., 2015). Strikingly, in both terminal boutons and elongated protrusions of Mem-G147W and Mem-K166del synapses, we observed either fragmentation of the normally continuous Futsch- and ANK2-XL-labeled cytoskeleton, or an absence of one or both proteins (Figure 6D). Thus, secretory defects due to Membrin mutations reduce the local integrity of the presynaptic cytoskeleton.

### Physiological abnormalities at Membrin mutant synapses

Finally, given the profound synaptic morphological abnormalities at Membrin mutant NMJs and cortical myoclonus and generalized epilepsy in *GOSR2*-PME patients, we asked whether Membrin mutations altered spontaneous or evoked neurotransmitter release at the L3 larval NMJ. We detected a clear reduction in the frequency of spontaneous miniature excitatory post-synaptic potentials (mEPSPs) in Mem-G147W and Mem-K166del (Figure 7A, B), while the amplitude and time course of mEPSPs were comparable between wild-type and the Membrin mutants (Figure S7A, B). No effect of Mem-G147W and Mem-K166del mutations on the amplitude of single postsynaptic evoked EPSPs was observed (Figure S7C). However, we often observed grossly deformed trains of EPSPs in both Mem-G147W and Mem-K166del following 5 consecutive stimuli at 10 Hz, where between one and all five EPSPs exhibited broader waveforms with multiple peaks and occasional merging of EPSPs (Figure 7C). Significantly more EPSP trains were abnormal in both mutants compared to Mem-WT: 5% of EPSP trains in Mem-WT were scored abnormal by a blinded observer, compared to ~ 25% in Mem-G147W and 22% in Mem-K166del (Figure 7D). In addition, the area under the EPSP train was robustly increased in Mem-G147W and Mem-K166del compared to Mem-WT (Figure 7E). These findings indicate that the morphological abnormalities caused by pathogenic *GOSR2* mutations are also associated with altered synaptic function.

**Figure 7.**
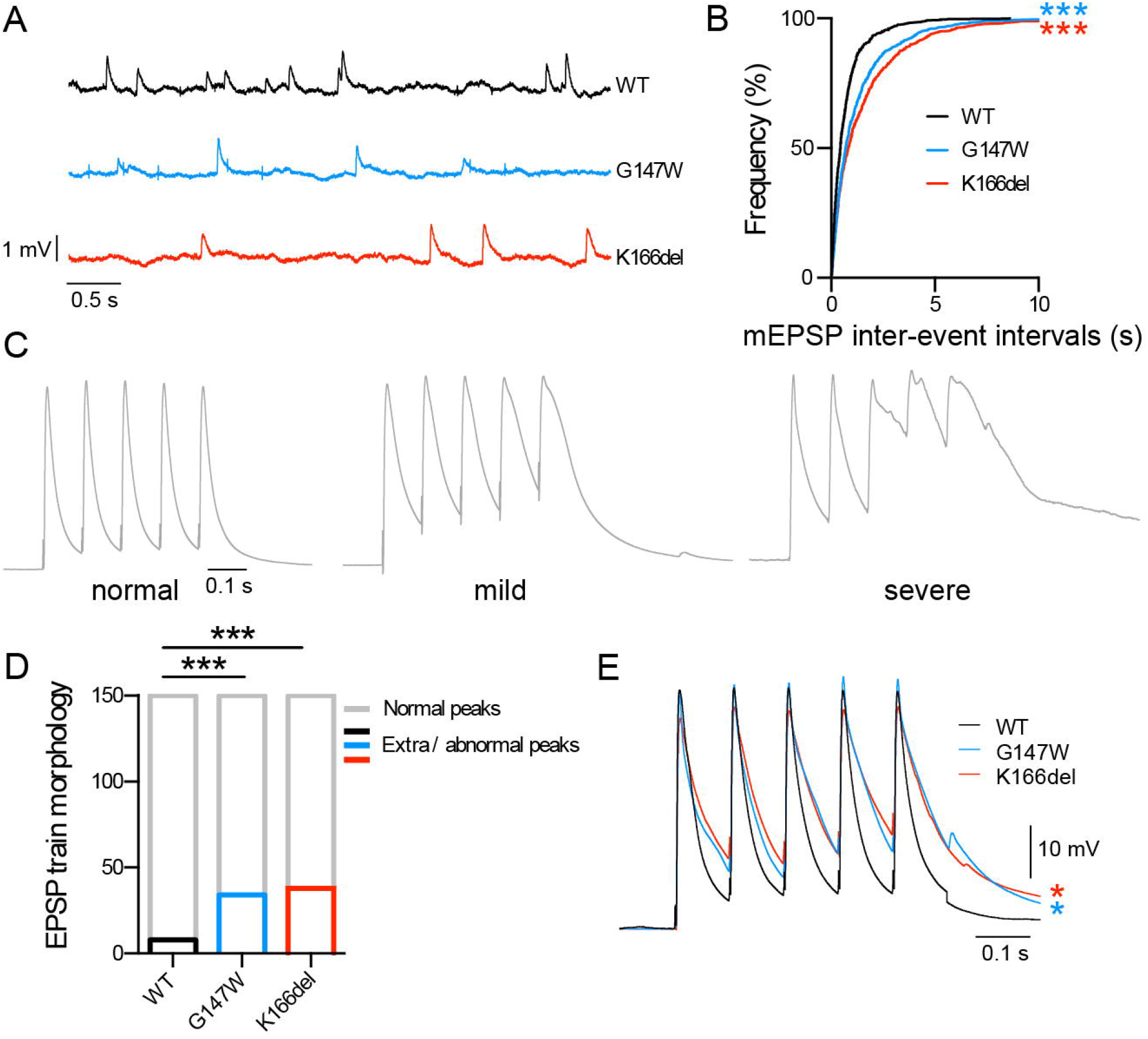
Physiological abnormalities at *membrin* mutant NMJs. (A) Representative traces of miniature excitatory post synaptic potentials (mEPSPs) recorded from Mem-WT/-G147W/-K166del L3 larval muscle 6 abdominal segments 2-4. (B) Cumulative frequency plot of mEPSP intervals. 800 events per genotype from 8 animals each are shown. (C) Illustrative traces depicting mild to severe EPSP waveform distortion following 5 stimuli at 10 Hz. Traces are normalized to the peak amplitude. (D) Analysis of total number of abnormal events as a result of 10 Hz stimulation. 15 events were analysed from each recording from 10 animals per genotype. (E) Overlay of averaged 10 Hz EPSP trains illustrating a significantly larger mean area under the curve in Mem-G147W and Mem-K166del compared to Mem-WT. n = 10. *, *** represents p < 0.05, 0.001 respectively; Kolmogorov-Smirnov test and Bonferroni correction (B), Fisher’s exact test and Bonferroni correction (D), one-way ANOVA with Dunnett’s multiple comparison test (E).

## Discussion

To date, how mutations in Membrin - a ubiquitous and essential Golgi SNARE protein - manifest as a disorder restricted to the nervous system has been unclear. Here we demonstrate that PME-causing Membrin mutations partially reduce SNARE activity yet still result in profound dendritic growth deficits in *Drosophila* models. Furthermore, we identify signatures of synaptic disassembly and altered neurotransmission, establishing a close dependence of synaptic stability and physiology upon precisely tuned secretory trafficking.

In *Drosophila*, the orthologous G147W and K166del Membrin mutations reduce dendritogenesis with a relative severity consistent with the corresponding effects on Bos1-dependent liposome fusion *in vitro*. Our findings reinforce the results of a *Drosophila* screen that identified the ER-to-Golgi trafficking proteins Sar1, Sec23 and Rab1 to be required for dendrite growth (Ye et al., 2007). Because our results are based on the study of a human Mendelian disorder, they provide an instance that highlights the importance of this pathway for human neuronal physiology and disease (Jan and Jan, 2010). Interestingly, mutations in Sec23A, Sec23A, Sec24D and Sar1b present in humans with largely non-neuronal clinical phenotypes of cranio-lenticulo-sutural dysplasia, congenital dyserythropoietic anemias, a syndromic form of osteogenesis imperfecta and lipid absorption disorders (Annesi et al., 2007; Bianchi et al., 2009; Boyadjiev et al., 2006; Garbes et al., 2015; Jones et al., 2003; Schwarz et al., 2009). This appears to be a consequence of tissue-specific differential utilization of the two available isoforms of Sec23 and Sar1 and special demands upon the COPII coat due to the large size of procollagen and chylomicrons (Canty and Kadler, 2005; Fromme et al., 2008; 2007; Jones et al., 2003).

We found that even subtle defects in the secretory pathway – such as observed in G176W Bos1/G144W Membrin – can be highly relevant for neuronal growth, while not critically affecting ER-to-Golgi trafficking in a non-neuronal cell type such as a fibroblast. We thereby provide experimental evidence for the postulate that, due to the unique plasma membrane demands of a growing neuron, even minor disruptions along the secretory pathway can selectively disrupt the nervous system while falling below a critical threshold in other organs (Pfenninger, 2009). We posit that this is the likeliest explanation of why *GOSR2*-PME is a syndrome restricted to the nervous system. Furthermore, the observed dendritic growth deficits might directly explain one hallmark of this disorder – lack of coordination. A dendritic growth bottleneck will presumably affect most severely neurons with highly elaborate dendritic arbors, such as cerebellar Purkinje cells (Ramón y Cajal, 1906). Impairment of these cells in turn would give rise to ataxia, as they are of critical importance for motor coordination (Cerminara et al., 2015; Kasumu and Bezprozvanny, 2010). Interestingly, cerebellar defects have even been suggested to also be involved in the pathogenesis of cortical myoclonus (Ganos et al., 2014).

Our study extends the findings of Ye et al. by showing that early secretory pathway changes can significantly impact synaptic morphology and physiology as well as dendritogenesis (Ye et al., 2007). We found that larval NMJs of Mem-G147W and Mem-K166del exhibit synaptic retraction, abnormal elongated protrusions lacking synaptic specializations, reduced spontaneous neurotransmitter release and malformed EPSPs. Interestingly, EMG evidence of motoneuron denervation in *GOSR2*-PME patients has been reported, and the typical absence of deep-tendon reflexes in this disorder might also be a consequence of such changes (van Egmond et al., 2014).

Membrin acts as a gatekeeper at the cis-Golgi and thereby likely determines the transition speeds for a plethora of synaptic and axonal proteins. Thus, the observed synaptic changes likely arise from a complex interaction of trafficking delays or steady-state reductions of many different proteins. Nevertheless, we identify several molecular correlates of the above structural changes. These include the loss or fragmentation of the cytoskeletal proteins ANK2-XL (an Ankyrin-2 isoform) and Futsch, particularly in presynaptic boutons with elongated protrusions. The axonal and synaptic cytoskeleton contains several interlinked constituents, including a microtubule core; actin filaments; and a submembranous mesh of Ankyrin and Spectrin (Goellner and Aberle, 2012). Collectively, these components regulate an array of key neurodevelopmental and physiological parameters including synaptic growth, morphology, and stability; axonal caliber; and ion channel localization (Chung et al., 2006; Jenkins and Bennett, 2001; Koch et al., 2008; Pan et al., 2006; Pielage et al., 2005; 2008; 2011; Roos et al., 2000; Stephan et al., 2015). Futsch is the *Drosophila* homologue of the mammalian microtubule-binding protein MAP1B (Roos et al., 2000), and the absence or fragmentation of Futsch in Mem-G147W and Mem-K166del synapses implies similar alterations in microtubule stability. Given the synaptic retraction observed in Mem-G147W and Mem-K166del NMJs, it is interesting to note that destabilization of microtubules is an early event during naturally occurring synapse loss at the mammalian NMJ (Bishop et al., 2004; Brill et al., 2016). Ankyrin-2 isoforms act to delineate *Drosophila* synaptic termini into characteristic rounded boutons separated by thin inter-bouton domains (Koch et al., 2008; Kurshan et al., 2009; Pielage et al., 2008). Since Mem-G147W and Mem-K166del boutons often exhibit elongated protrusions reminiscent of extended inter-bouton domains, we speculate a direct link between local destabilization of the synaptic cytoskeleton, synaptic retraction and the presence of elongated protrusions in Mem-G147W and Mem-K166del synapses.

Membrin mutations not only affect synaptic structure but also disrupt evoked and spontaneous neurotransmitter release. We can only speculate about the underlying mechanisms, which are likely to again represent a complex, cumulative process caused by insufficient trafficking of diverse cargos such as ion channels and proteins required for presynaptic vesicle fusion. Increased variability of axonal diameters in Mem-G147W and Mem-K166del may further alter action potential propagation velocities across different axons innervating the same muscle, leading to de-synchronized depolarizing currents.

The progressive myoclonus epilepsies are phenotypically heterogeneous, with several different causative genes known to date (Michelucci et al., 2012). Some of the PMEs with more closely related phenotypes might share cellular pathways and/or neural circuits with *GOSR2*-PME. For instance, mutations in *PRICKLE1* cause PME in humans and seizures in flies (Bassuk et al., 2008; Tao et al., 2011), and *PRICKLE1* has been linked to neurite growth and axonal trafficking (Ehaideb et al., 2014; Liu et al., 2013). PME-linked mutations in the potassium channel gene *KCNC1* are thought to mainly impair fast-spiking neurons (Muona et al., 2015; Oliver et al., 2017). Such a preferential defect in high-frequency firing neurons is also conceivable in *GOSR2*-PME, where EPSP dys-morphologies are more pronounced under repetitive stimulation.

In summary, by elucidating the pathophysiology of *GOSR2*-PME we identify a critical role for Membrin in promoting synaptic integrity, highlight stringent requirements of dendritic growth on the secretory pathway, and define how mutations in an essential gene can selectively disrupt nervous system function.

## Materials and Methods

### Molecular biology

Human *GOSR2* CDS as well as *Drosophila melanogaster membrin* CDS with and without the G144W/G147W and K164del/K166del mutations preceded by 5’ FLAG tag coding sequence were custom synthesized by GeneArt (Thermo Fisher Scientific) and subsequently cloned via NotI and KpnI (NEB) into pUASTattB, giving rise to pUASTattB_FLAG::*GOSR2*[WT]/pUASTattB_FLAG::*membrin*[WT], pUASTattB_FLAG::*GOSR2*[G144W]/pUASTattB_FLAG::*membrin*[G147W] and pUASTattB_FLAG::*GOSR2*[K164del]/pUASTattB_FLAG::*membrin*[K166del]. The *GOSR2* constructs were also cloned into of pcDNA3.1(−) by replacing the CDS of pcDNA3.1(−) mouse C/EBP beta (LAP) (Addgene plasmid #12557) for these inserts via NotI and KpnI (NEB), giving rise to pcDNA3.1(−)_*GOSR2*[WT], pcDNA3.1(−)_*GOSR2* [G144W] and pcDNA3.1(−)_*GOSR2*[K164del]. pGEX-2T-GST::*Sed5*, pET28a-His::*Sec22*, pET28a-His::*Bet1* and pET28a-His::*Bos1*[WT] were previously described (Parlati et al., 2000; 2002). To generate pET28a-His::*Bos1*[G176W] and pET28a-His::*Bos1*[D196del], the respective mutations were introduced with the QuickChange Site-Directed Mutagenesis Kit (Agilent Technology). The Halo::FM4::hGH containing plasmid was previously cloned in the Rothman lab (Lavieu et al., 2013).

### Bioinformatics

SNARE motifs in human and *Drosophila* Membrin, as well as yeast Bos1, were identified as described previously (Kloepper et al., 2007) and aligned with Clustal Omega (McWilliam et al., 2013). Conserved residues were highlighted using BoxShade.

### Liposome fusion assays

The recombinant yeast Golgi SNARE proteins were expressed and purified in *E. coli* BL21 (DE3) cells as described previously (Parlati et al., 2000). GST-Sed5 was purified using glutathione agarose affinity beads (Thermo Fisher Scientific) and the GST-tag was removed using 100U of human thrombin (Sigma) with overnight incubation at 4°C in Buffer A (25 mM HEPES pH7.4, 400 mM KCl, 10% glycerol, 1 mM DTT, 1% n-Octyl-β-D-glucopyranoside) containing 2 mM CaCl_2_. The other proteins – Sec22, Bet1, wild-type and G176W/D196del mutant Bos1 – were purified with a His^6^-tag using HisPur Ni-NTA affinity beads (Thermo Fisher Scientific) in Buffer A containing 300 mM imidazole, pH 7.5.

Purified SNARE proteins were reconstituted into lipid vesicles using the detergent (1% n-Octyl-β-D-glucopyranoside) dilution and dialysis method (Weber et al., 1998). For the t-SNARE acceptor liposomes, 27 μM of Sed5, Sec22 and wild-type or G176W/D196del mutant Bos1 in 500 μl were incubated overnight at 4°C and then incorporated into palmitoyl-2-oleoyl phosphatidylcholine (POPC): 1,2 dioleoyl phosphatidylserine (DOPS) at 85:15 mol% liposomes. The donor liposomes containing Bet1 (1:100 protein:lipid ratio) were prepared similarly with the lipid mix of POPC, DOPS and the fluorescent probes Nitro-2-1,3 benzoxadiazol-4yl-phosphatidylethanolamine (NBD-PE) and Rhodamine-PE (1.5 mol% each). All lipids were purchased from Avanti Polar lipids (Alabaster, AL). Liposome fusion assay was performed by mixing 5 μl of the donor liposome (Bet1) with 45 μl of the acceptor liposome (t-SNAREs). Fusion of liposomes was monitored by the change in NBD fluorescence at 538 nm using a Flexstation 3 microplate reader (Molecular Devices). After 120 min, 10 μl of 5% w/v *n*-dodecyl-β-maltoside (Thermo Fisher Scientific) was added to lyse all vesicles to estimate the maximum NBD fluorescence (Weber et al., 1998). For experiments with the Bet1 peptide, the t-SNARE liposomes were pre-incubated at 37°C for 45 min with 50 μM peptide corresponding to the C-terminal half of Bet1 (RGSNQTIDQLGDTFHNTSVKLKRTFGNMMEMA, Vc peptide) prior to addition of Bet1 liposomes to initiate fusion.

### Cell culture and transfections

Primary skin derived fibroblasts from the first described *GOSR2*-PME patient were kindly shared by Mark Corbett (Corbett et al., 2011). As controls, we used fibroblasts from healthy individuals of either the same sex and similar age, or opposing sex and divergent age (control 1 = 23 year old female; control 2 = 60 year old male at time of biopsy). Fibroblast and HEK293T cells were grown in DMEM + 10% FBS at 37°C and 5% CO_2_. Fibroblast transfections were carried out with lipofectamine 2000 (Thermo Fisher Scientific), HEK293T transfections with Effectene (Qiagen).

### Fibroblast imaging

For immuno-fluorescence studies cells were seeded on #1.5 glass coverslips, fixed with 4% PFA and permeabilized in PBS containing Triton-X100 and NP40. After primary and secondary antibody incubation steps coverslips were mounted in SlowFade Gold Antifade (Thermo Fisher Scientific). The following antibodies were used: mouse anti-Membrin (clone 25, BD Biosciences; This antibody was raised against Membrin residues 5-124 and therefore should not be affected by the G144W mutation.), mouse anti-FLAG (M2 clone, Sigma), rat anti-FLAG (Agilent), rabbit anti-GPP130 (Cambridge Bioscience), rabbit anti-PDI (Sigma), and goat anti-mouse/rabbit/rat Alexa Fluor 488/555/647 conjugated secondaries (Thermo Fisher Scientific). For Golgi trafficking studies cells were loaded with HaloTag TMR Ligand (Promega) 24 h post transfection with Halo::FM4::hGH. Subsequently ER retained cargo was realeased by addition of D/D solubilizer 1.5 μM (Clontech). Fixed samples were imaged with a Plan-Apochromat 63x 1.4 NA oil immersion objective on Zeiss confocal LSM710 or LSM880 microscopes.

### Western blot

Cells or whole L3 larvae were lysed in 20 mM HEPES pH 7.5, 100 mM KCl, 5% glycerol, 10 mM EDTA, 1% Triton X-100 supplemented with phosphatase and proteinase inhibitors (PhosSTOP/cOmplete, Roche). Total protein content was quantified with the Pierce 660 nm assay (Thermo Fisher Scientific) and equal amounts loaded into each lane. Proteins were separated on a 4-10% Bis-Tris polyacrylamide gel (Thermo Fisher Scientific) and transferred onto PVDF membranes (EMD Millipore). The following antibodies were used: mouse anti-Membrin (clone 25, BD Biosciences), mouse anti-β-actin (clone AC-74, Sigma) and HRP-conjugated anti-mouse (Jackson Immuno). For semi-quantitative western blots, detection was carried out with SuperSignal West Pico Chemiluminescent Substrate (Thermo Fisher Scientific) and a ChemiDocTM Imaging system (Bio-Rad). Band intensities were extracted with Image Studio Lite (Li-cor).

### *Drosophila* stocks

*membrin*^1524^ flies were previously generated in an EMS screen and kindly shared by Mark Krasnow (Ghabrial et al., 2011). This strain harbors a premature stop codon upstream of the *membrin* SNARE domain encoding sequence and therefore represents a null-allele. To control for potential genetic background effects we outcrossed *membrin*^1524^ for five generations into an isogenic iso31 background by following an AccI (NEB) restriction site that is introduced by the nonsense mutation. *GOSR2* and *membrin* transgenic flies were generated by microinjection of pUASTattB_FLAG::*GOSR2*[WT/G144W/K164del]/pUASTattB_FLAG::*membrin*[WT/G147 W/K166del] into *y*[1] M{vas-int.Dm}ZH-2A w*; M{3xP3-RFP.attP’}ZH-51C embryos (Cambridge fly facility). This approach enabled us to retrieve wild-type and mutant *membrin* transgenic fly lines with insertions into precisely the same genomic locus (ZH-51C), which is important to provide comparable transgene expression levels (Bischof et al., 2007). In order to express wild-type or mutant *GOSR2* or *membrin* in a *membrin* null background the following stocks were created and crossed to each other: I. *w*[1118]; UAS-FLAG::*GOSR2*[WT/G144W/K164del]; *membrin*^1524^/TM6B, *tb* or *w*[1118]; UAS-FLAG::*membrin*[WT/G147W/K166del]; *membrin*^1524^/TM6B, *tb* and II. *w*[1118]; +; *membrin*^1524^, daughterless-GAL4/TM6B, *tb*. Each component of these flies was outcrossed for five generations into iso31 prior to assembly by standard mating schemes. For dendritic analysis, *ppk*-CD4::tdGFP (Han et al., 2011) was incorporated into stock II. (*w*[1118]; *ppk*-CD4::tdGFP; *membrin*^1524^ *daughterless*-Gal4/TM6B, *tb*) and crossed to stock I. *daughterless*-Gal4 was obtained from Bloomington stock center. Flies were reared on a standard cornmeal-molasses-yeast medium at 25°C in 12 h light-dark cycle.

### *Drosophila* viability and locomotion

To assess viability of *membrin* mutant animals, the above I x II crosses were allowed to egg-lay onto apple-juice agar plates overnight. Subsequently, eggs/embryos were counted and transferred to standard food tubes. After completion of pupation, non-*tubby* pupae were counted. Given that theoretically only one quarter of the collected egg/embryos are of the correct genotype, we used the following formula to calculate egg/embryo to pupa viability: non-*tubby* pupae/(total eggs/4). The resulting fraction was normalized to wild-type because we found a considerable reduction with consecutive egg-lays, presumably reflecting decreasing fertilization. Eclosion rates were determined 11 days after onset of egg laying, a time point where under our conditions non-eclosion was equal to death or imminent death in the pupal case. To assess locomotion, L3 larvae were placed in the center of a 100 mm sucrose-agar plate positioned on top of a 4 mm grid, allowed to settle for 30 s, and filmed for the next 60 s. Grid-breaks in this time period were then assessed offline.

### Dendritic analysis

The highly elaborate ddaC neuron in abdominal segment 5 of L3 larvae was used throughout (Grueber et al., 2002). For morphological analysis, larvae were heat-killed and mounted under a #1.5 glass coverslip. Z-stacks of ddaC neurons were obtained with Zeiss confocal LSM710 microscopes with a N-Achroplan 10x 0.25 NA objective to capture the entire arbor. To extract total dendrite length and to serve as a template for the ImageJ Sholl Analysis plugin, dendrites were semi-manually traced with the ImageJ NeuronJ plugin (Ferreira et al., 2014; Meijering et al., 2004). Terminal branches were manually counted on dendrite tracings with the ImageJ multi-point tool. For FRAP experiments, L3 larvae were fillet-prepped in HL3 saline without Ca^2+^ (70 mM NaCl, 5 mM KCl, 20 mM MgCl2, 10 mM NaHCO3, 5 mM trehalose, 115 mM sucrose, 5 mM HEPES, pH 7.2). Fillets were transferred to a #1.5 glass bottom dished and submerged in fresh HL3 saline with a custom-made platinum wire anchor before being imaged on an inverted Zeiss confocal LSM510 with a Plan-Apochromat 20x 0.8 NA objective. Bleaching of a 50 μm^2^ area encompassing major primary dendrites directly adjacent to the soma was carried out by scanning for 200 iterations with 100% 488 nm transmission. Mean fluorescence intensity of a small dendritic region contained in a 5 μm circle 25 μm from the bleach border adjacent to the soma served as a read-out. This is the most distant dendritic region from either bleach margin and thus the contribution to fluorescence recovery from lateral diffusion of dendrite surface localizing CD4::tdGFP is minimized. Bleach depth in this region was consistently greater than 87%.

### Immuno-histochemistry of larval neuromuscular junctions and brains

When examining synaptic development at the larval neuromuscular junction (NMJ), synapses innervating muscle 6/7 of segment 3 were imaged on a Zeiss confocal LSM710 with either a Plan-Apochromat 20x 0.8 NA or a Plan-Apochromat 63x 1.4 NA oil immersion objective. Late-stage L3 larvae were dissected in low Ca^2+^ (0.2 mM) HL3 (see above). Dissected NMJ preparations were fixed in either 4% PFA or Bouin’s solution for 10-20 min at room temperature. Larval brains were dissected and immunostained as described previously (Wu and Luo, 2006). The following antibodies were used: mouse anti-BRP (clone nc82, DSHB), anti-CSP (clone 6D6, DSHB), anti-GLURIIA (clone 8B4D2, DSHB), and anti-Futsch (clone 22C10, DSHB); rabbit anti-ANK2-XL (kind gift from Herman Aberle), rabbit anti-SNAP-25 (kind gift from David Deitcher), rabbit anti-GLURIII and rabbit anti-vGLUT (kind gifts from Aaron DiAntonio), mouse anti-GFP (clone 3E6, Thermo Fisher Scientific), goat anti-HRP Alexa Fluor 488 conjugated (Jackson ImmunoResearch) and secondaries as above. For quantification of synaptic development, all experiments were performed and analysed blind to experimental genotype. Boutons were identified using anti-HRP and anti-CSP (Zinsmaier et al., 1990). Both type 1b and type 1s boutons were included in the bouton count. The number of BRP puncta and CSP-positive boutons was quantified using the ImageJ 3D Object Counter Plugin (Bolte and Cordelières, 2006). Since CSP levels were reduced in GOSR2-PME model backgrounds, CSP fluorescent signals were enhanced to saturating levels in all genotypes prior to bouton counting.

### NMJ electrophysiology

Wandering L3 larvae were dissected in ice-cold, Ca^2+^-free HL3.1-like solution (70 mM NaCl, 5 mM KCl, 10 mM NaHCO_3_, 115 mM sucrose, 5 mM trehalose, 5 mM HEPES, 10 mM MgCl_2_). Motor nerves were severed just below the ventral nerve cord and the brain was removed. CaCl_2_ (1 mM) was added to the bath solution for intracellular recording from muscle 6 of abdominal segments 2-4. Sharp microelectrodes (thick-walled borosilicate glass capillaries, pulled on a Sutter Flaming/Brown P-97 micropipette puller) were filled with 3M KCl and had resistances of 20 – 30 MΩ. For recording of stimulus evoked excitatory post synaptic potentials (EPSPs), severed nerves were drawn into a thin-walled glass-stimulating pipette and stimulated with square-wave voltage pulses (0.1 ms, 10 V, A-M Systems Model 2100 Isolated Pulse Simulator). EPSPs and spontaneously-occurring miniature EPSPs (mEPSPs) were recorded at a controlled room temperature of 22-25°C with a Geneclamp 500 amplifier (Axon Instruments) and were further amplified with a LHBF-48x amplifier (NPI Electronic). The membrane potential was set to −70 mV with current injection at the start of each recording. Voltage signals were low-pass filtered at 1.67 kHz (10 kHz 4 pole Bessel on Geneclamp 500, 1.7 kHz 8-pole Bessel on LHBF-48x) and digitised at 25 kHz by a CED-1401 plus A/D interface (Cambridge Electronic Design, UK) using Spike2 software (v. 5.13) (CED, Cambridge, UK). Synaptic potentials were analysed offline using Strathclyde Electrophysiology Software WinEDR (v3.5.2) and GraphPad Prism (v.6). All synaptic events were verified manually. Recordings were discarded if the initial resting membrane potential was more positive than −60 mV or varied by more than 10% throughout the recording. mEPSPs were recorded for a minimum of 5 min. Single EPSPs were evoked 10 times at a standard frequency of 0.033 Hz or trains of 5 EPSPs at 10 Hz were evoked 3 times at 0.033 Hz. Intervals and amplitudes of mEPSPs were compared by creating a cumulative distribution for each genotype of 800 measurements across 8 animals, with each animal contributing 100 values. To analyse the mEPSP waveform, a mean mEPSP was constructed for each recording from events that showed only a single clear peak and a smooth decay so as to prevent distortion of the waveform by closely-occurring mEPSPs.

## Author Contributions

Conceptualization, R.P., S.S.K, J.E.R. and J.E.C.J.; Methodology, R.P., N.T.M., S.S.K. and J.E.C.J.; Formal Analysis, R.P., S.A.L., N.T.M., J.E.C.J.; Investigation, R.P., S.A.L., N.T.M., J.E.C.J.; Resources, H.H., D.M.K., M.M.U., J.J.L.H., J.E.R. and J.E.C.J.; Writing – Original Draft, R.P. and J.E.C.J.; Writing – Review & Editing, R.P., S.A.L., N.T.M., D.M.K., M.M.U., S.S.K., J.J.L.H., J.E.C.J.; Visualisation, R.P. and J.E.C.J.; Supervision, H.H., D.M.K., M.M.U., S.S.K., J.J.L.H., J.E.R., J.E.C.J.; Project Administration, R.P. and J.E.C.J.; Funding Acquisition, R.P., H.H., D.M.K., M.M.U., J.J.L.H., J.E.R., J.E.C.J.

## Acknowledgements

We would like to thank Andreas Ernst for experimental suggestions and discussion of data, Mark Corbett for kindly sharing G144W-Membrin mutant fibroblasts, Mark Krasnow for providing *membrin*^1524^ flies, Lily Yeh Jan and Yuh-Nung Jan for sharing *ppk*>CD4::tdGFP flies and Herman Aberle, David Deitcher and Aaron DiAntonio for sharing antibodies, and Giampietro Schiavo, Helene Plun-Favreau and Stephanie Schorge for helpful comments on the manuscript. This work was funded by a Wellcome Trust Strategic Award (J.E.C.J, J.E.R, H.H and D.M.K), the MRC (MR/P012256/1: J.E.C.J), the BBSRC (BB/J017221/1: J.J.L.H.) and NIHR funding for UCL/UCLH Biomedical Research Centre (BRC). R.P. is supported by a Brain Research Trust PhD studentship. S.A.L is supported by a BBSRC DTP studentship.

